# Characterisation of neonatal *Staphylococcus capitis* NRCS-A isolates compared with non NRCS-A *Staphylococcus capitis* from neonates and adults

**DOI:** 10.1101/2022.12.02.518718

**Authors:** Heather Felgate, Dheeraj Sethi, Kirstin Faust, Cemsid Kiy, Christoph Härtel, Jan Rupp, Rebecca Clifford, Rachel Dean, Catherine Tremlett, John Wain, Gemma Langridge, Paul Clarke, Andrew Page, Mark A Webber

## Abstract

*Staphylococcus capitis* is a frequent cause of Late-Onset Sepsis (LOS) in neonates admitted to Neonatal Intensive Care Units (NICU). The NRCS-A clone of *S. capitis* has been isolated from NICUs globally although the reasons for the global success of this clone are not understood.

We analysed a collection of *S. capitis* colonising babies admitted to two NICUs, one in the UK and one in Germany as well as corresponding pathological clinical isolates. Genome analysis identified 3 groups; non-NRCS-A isolates, NRCS-A isolates, and a group of ‘proto NRCS-A’ - isolates closely related to NRCS-A but not associated with neonatal infection. All bloodstream isolates belonged to the NRCS-A group and were indistinguishable from strains carried on the skin or in the gut. NRCS-A isolates showed increased tolerance to chlorhexidine and antibiotics relative to the other *S. capitis* as well as enhanced ability to grow at higher pH values. Analysis of 138 pangenomes of the clades identified characteristic *nsr* and *tarJ* genes in the NRCS-A and proto groups with a CRISPR-cas system only seen in NRCS-A isolates which also showed enrichment of genes for metal acquisition and transport.

We found evidence for transmission of *S. capitis* NRCS-A within NICU, with related isolates shared between babies and multiple acquisitions by some babies. Our data show NRCS-A strains commonly colonise uninfected babies in NICU representing a potential reservoir for potential infection. This work provides more evidence that adaptation to survive in the gut and on skin facilitates spread of NRCS-A, and that metal acquisition and tolerance may be important to the biology of NRCS-A. Understanding how NRCS-A survives in NICUs can help develop infection control procedures against this clone.

## Introduction

Non-*aureus* Staphylococci are common commensal bacteria, however many have also been implicated in nosocomial infections and are also found to cause Late Onset Sepsis (LOS). LOS is defined as sepsis occurring ≥ 72 hrs after birth which can be caused by various bacterial species, the most common being the Non-*aureus* Staphylococci (NAS) (Bentlin et al., 2010, Makhoul et al., 2005, Dong and Speer, 2014, Harvey et al., 2022). LOS increases the length of hospital stay, invasive procedures, and provokes more and longer antibiotic treatments all of which can have negative impacts on the long-term outcomes of new-born babies (Bentlin et al., 2010, Makhoul et al., 2005, Dong and Speer, 2014, Rasigade et al., 2012).

*Staphylococcus capitis* is a common commensal of humans and is regularly cultured from skin or nasal swabs (Becker et al., 2020), but a particular clone of *S. capitis* NRCS-A has been found to associated with LOS. Originally identified in France (the archetypal strain is known as CR01) it has subsequently been found in neonatal intensive care units (NICUs) in 17 countries (Lemriss et al., 2014, Lepainteur et al., 2013, Wirth et al., 2020, Butin et al., 2017, Harvey et al., 2022). Infection with NRCS-A has been associated with a higher morbidity and mortality than other NAS (Butin et al., 2019, Ben Said et al., 2016). The transmission routes of NRCS-A are not well understood but it has been documented to colonise incubators and equipment and has reduced susceptibility to antiseptics which has been proposed to aid its environmental survival. Unlike other NAS, infection with NRCS-A is not associated with maternal-fetal transmission, and babies born by Caesarean-section have been shown to be colonised more often than those delivered vaginally (Butin et al., 2019).

NRCS-A isolates typically have a multidrug resistance profile including methicillin, aminoglycosides and fosfomycin, hetero-resistance to vancomycin (a common treatment for LOS) and decreased susceptibility to chlorhexidine (Wirth et al., 2020, Lepainteur et al., 2013, Rasigade et al., 2012, Harvey et al., 2022). Many NRCS-A isolates have been found to contain a *SCCmec* mobile genetic element (*SCCmec-SCCcad-SCCars-SCCcop*) responsible for the methicillin resistance (Carter et al., 2018, Du et al., 2021, Becker et al., 2014) and decreased susceptibility to vancomycin has been associated with SNPs in *sarA* and *glnQ* (Wirth et al., 2020). Analysis of genomes of a panel of NRCS-A isolates from various sites and sources suggested the clone emerged in the 1960s before expanding in the 1980s with vancomycin use proposed as a major driver in the selection and evolution of NRCS-A (Wirth et al., 2020).

*S. capitis* infection has only been studied sporadically outside of the neonatal setting but *S. capitis* has been identified in bone and joint infections and, whilst rarely associated with adult disease, NRCS-A has been identified in endocarditis, osteomyelitis and prosthetic joint infections (PJI) in adults (Tevell et al., 2020).

Here we report results from a longitudinal survey of NAS from skin and gut swabs taken from babies on NICUs from two countries (the UK and Germany) where different antiseptic regimens were in place (Sethi et al., 2021). Over 1000 NAS isolates were characterised for antimicrobial susceptibility and sequenced. From this collection we have analysed the population structure of *S. capitis* and compared carriage isolates with those from neonatal blood cultures as well as additional isolates from healthy adult skin swabs and PJI.

We found that NRCS-A strains are commonly carried by uninfected neonates representing a reservoir of potential infection and identified probable transmission between babies on NICU. We also identified a clade closely related to NRCS-A but without an association with disease. Comparison of the NRCS-A isolates with this clade (‘proto NRCS-A’) identified genomic and phenotypic features which may contribute to the success and prevalence of this epidemic clone.

## Methods

### Collection of isolates of *S. capitis*

Infants admitted to the NICUs of the Norfolk and Norwich University Hospital (Norwich, UK) or University Children’s Hospital (Lubeck, German) over 10-week periods in 2017 or 2018 were enrolled in this study as recently described (Sethi et al., 2021). Swabs are taken routinely from babies upon admission and throughout their stays in both sites for MRSA surveillance; duplicate swabs were taken for this study and staphylococci isolated. In addition, isolates from positive blood, cerebrospinal fluid, urine, and wound cultures were included, if taken during the study period.

The UK unit enrolment was between November 2017 and January 2018, and the German enrolment was from January to March 2018. Amies Charcoal Swabs (Thermo Fisher Scientific) were used to isolate bacteria from all infants on admission and weekly until discharge. Swabs from the ear, nose, axilla, groin and gut were taken and streaked on 5% horse blood agar (Thermo Fisher), prior to incubation at 37 °C for 24 hours and final identification of coagulase-negative Staphylococci after sub-culture on mannitol-salt agar (Oxoid, Thermo Fisher Scientific), coagulase testing (MERCK; 75832), and/or MALDI-TOF mass spectrometry (Bruker) as previously described (Sethi et al., 2021). Isolates were stored on preservation beads at −80 ^o^C (Protect, Technical Service Consultants Ltd) and in 96 deep-well plates in 20% glycerol at −40 ^o^C. Clinical isolates of *S. copitis* from both units which were identified by local microbiology departments during the study periods as part of usual practice were also included as well as further anonymised clinical isolates taken from routine blood tests taken from neonates with suspected sepsis at the NNUH NICU collected in 2018 (n=7) and between June and May 2022 (n=5). In addition to isolates from neonates, a further panel of 15 *S. capitis* were taken from a pre-existing Staphylococci collection isolated from adult healthy skin swabs, and clinical isolates taken from adult blood cultures (where infection was suspected) and 5 recovered from PJI (Felgate et al., 2021).

### DNA extraction

Isolates were grown in 1 mL Brain Heart Infusion (BHI, MERCK) broth overnight at 37 °C. Cultures were pelleted and resuspended in 100 μl 0.5 mg/mL lysostaphin (from *Staphylococcus staphylolyticus*, MERCK) and incubated at 37 °C for a minimum of 1 hour. For Illumina sequencing DNA was extracted from the lysate with the Quick-DNA Fungal/Bacterial 96 kit (Cambridge Bioscience), in accordance with the manufacturer’s guidelines. DNA was quantified using the Quant-iT^™^ dsDNA HS assay (ThermoFisher), and fluorescence was measured on a FLUOstar Optima plate reader at 480/530 nm (excitation/emission).

For high molecular weight DNA, the Quick-DNA HMW MagBead Kit (Zymo) was used with a modified protocol. Starting culture volume was increased to 500 μl and 50 μl of 0.5 mg/mL lysostaphin (from *Staphylococcus staphylolyticus*, MERCK) was added, instead of lysozyme. Isolates were incubated at 37 °C for 1 hour prior to adding ProteinaseK. DNA was quantified with dsDNA HS Qubit Assay (ThermoFisher).

### Sequencing

For Illumina sequencing genomic DNA was normalised to 5ng/μl in 10mM Tris-HCl. 0.5 μl of TB1 tagmentation DNA Buffer was mixed with 0.5 μl BLT, Tagment DNA Enzyme (Illumina) and 4 μl PCR grade water in a master mix and 5 μl added to a chilled 96 well plate. 2 μl of normalised DNA (10 ng total) was pipette mixed with the 5 μl of the tagmentation mix and heated to 55 ^o^C for 15 minutes in a PCR block. A PCR master mix was made up using 4 ul kapa2G buffer, 0.4 μl dNTP’s, 0.08 μl Polymerase and 4.52 μl PCR grade water, contained in the Kap2G Robust PCR kit (MERCK) per sample and 9 μl added to each well need to be used in a 96-well plate. 2 μl of each P7 and P5 of Nextera XT Index Kit v2 index primers (Illumina) were added to each well. Finally, the 7 μl of Tagmentation mix was added and mixed. The PCR was run with 72°C for 3 minutes, 95°C for 1 minute, 14 cycles of 95°C for 10s, 55°C for 20s and 72°C for 3 minutes. Following the PCR reaction, the libraries were quantified using the Promega QuantiFluor^®^ dsDNA System and run on a GloMax^®^ Discover Microplate Reader. Libraries were pooled following quantification in equal quantities. The final pool was double-SPRI size selected between 0.5 and 0.7X bead volumes using KAPA Pure Beads (Roche). The final pool was quantified on a Qubit 3.0 instrument and run on a D5000 ScreenTape (Agilent) using the Agilent Tapestation 4200 to calculate the final library pool molarity.

The pool was run at a final concentration of 1.5 pM on an Illumina Nextseq500 instrument using a Mid Output Flowcell (NSQ^®^ 500 Mid Output KT v2(300 CYS) Illumina) following the Illumina recommended denaturation and loading recommendations which included a 1% PhiX spike in (PhiX Control v3 Illumina). Data was uploaded to Basespace (www.basespace.illumina.com) where the raw data was converted to 8 FASTQ files for each sample.

For long read sequencing up to 400 ng of DNA was incubated with Ultra II End-prep reaction buffer and Ultra II End-prep enzyme mix for end repair. The samples then had native barcodes (NB01-24) added with the addition of the NEB Blunt/TA Ligase Master Mix and then the samples were heated on thermal cycler at 20 °C for 20 m, 65 °C for 10 m. The libraries were pooled and cleaned using AMPure XP beads, DNA was quantified using dsDNA HS Qubit (ThermoFisher) and Tapestation (Aligent). For the adaptor ligation step, Adapter Mix II (AMX II), NEBNext Quick Ligation Reaction Buffer (5X) and Quick T4 DNA Ligase was added to the pooled library and cleaned using AMPure XP beads. The library was eluted and loaded on to a primed MinION flow cell (Oxford Nanopore Technologies). Base calling (June 2022) was carried out using Guppy (v6.06) (Oxford Nanopore Technologies).

### Genomic and Phylogenetic analysis

Once generated, sequence data was processed via a series of pipelines on a Galaxy instance hosted at the Quadram Institute Bioscience. FASTQ files were used to assigned a microbial classification and check for contamination for each sample using Centrifuge (v0.15) (Kim et al., 2016). *S. capitis* strains were then used for further analysis, isolates were assembled using SPAdes (v3.12.0) (Bankevich et al., 2012), with default parameters applied, and resulting assemblies analysed for quality with QUAST (v5.0.2) (Mikheenko et al., 2018). Isolates with less than X10 genomic coverage were omitted from the final collection.

Long read data was assembled using Flye (v2.5) (Lin et al., 2016, Kolmogorov et al., 2019) the Illumina short reads were then mapped to the Flye scaffolds using minimap2 (v2.17) (Li and Durbin, 2010, Li and Durbin, 2009) and the sequence files were then polished with 2 rounds of pilon (v1.20.1) (Walker et al., 2014).

The phylogeny of the *S. capitis* isolates was determined using gene presence/absence after creating a tree based on the core gene alignment. Prokka (v1.14.5) (Seemann, 2014) was used to annotate the assembled contigs along with nine available *S. capitis* reference genomes (https://www.ncbi.nlm.nih.gov, Supplementary Data 1). The GFF3 files were then submitted to ROARY (v3.13.0) to determine the core and accessory genomes (requiring that 75% percent of isolates should carry a gene to be considered to be in the core) (Page et al., 2015). A phylogenetic tree was inferred using IQTREE (v 1.6.12) (Nguyen et al., 2015) from the core gene alignment output from Roary, and then visualised and annotated in iTOL (Letunic I. and Bork, 2019).

The presence of antimicrobial resistance and virulence genes were identified using ABRicate (v2.13.2, with 75% minimum identity, and 90% minimum coverage filters applied) (Seeman, 2016) and ARIBA (Hunt et al., 2017) using the CARD database (v3.1.1) (Alcock et al., 2020).

To identify SNPs, FASTQ files were processed by Snippy (v4.4.3), and Snippy-core (v4.4.3) used to generate a core alignment FASTA which was then analysed by snp-dist (v0.6.3) (Seemann, 2015, Seeman, 2019). The reference genome used for all Snippy tools was the CR05 genome (NZ_CTEO01000001.1). The Snippy-core alignment was then used to create a phylogeny based on core SNPs and run through IQTREE (Nguyen et al., 2015), the resulting maximum likelihood tree was visualised using iTOL (Letunic I. and Bork, 2019).

Homologues of specific sequences of interest were identified using BLASTn, sequence data for the *SCCmec-SCCcad-SCCars-SCCcop* (KF049201.1) mobile element was used as a reference to identify other similar sequences in the collection using BLASTn megablast (v2.10.1; default settings)) (Camacho et al., 2009, Cock et al., 2015). The NCBI protein database was searched for homologues to Nsr (CDI72769.1) and TarJ (CDI72761.1) with BLASTp (Cock et al., 2015).

Alignment of CRISPR-Cas Type-III-A genes from different strains was assessed by taking a reference sequence from *S. capitis* CR05 and creating a Snippy-core full alignment which was then submitted to ClustalW (v1.91) (Larkin et al., 2007).

Long read sequences were compared using BLASTn pairwise alignment (NCBI, 2022) and visualised using Artemis Comparison Tool (Carver et al., 2008).

### Antimicrobial Susceptibility Testing

The susceptibility of all *S. capitis* isolates to the biocides chlorhexidine gluconate (CHX) and octenidine hydrochloride (OCT) and the antibiotics, benzylpenicillin (PEN), cefotaxime (CEF), daptomycin (DAP), gentamicin (GEN), fusidic Acid (FUS) and vancomycin (VAN) was determined in Mueller-Hinton agar (for daptomycin, Ca^2+^ was added to a final of concentration 50 mg/L) (Sigma-Aldrich) in accordance with EUCAST guidelines. Plates were inoculated with ~10^4^ cells diluted from overnight cultures grown in Mueller-Hinton broth (Sigma-Aldrich) using a multi-point inoculator (Denley). *Staphylococcus aureus* controls were run on all the plates, ST 239 (‘TW20’) and NCTC 8532 (‘F77’) were used on the biocide plates and ATCC 29213 for the antibiotic plates. The plates were incubated at 37 ^o^C for 24 hours before minimum inhibitory concentrations (MICs) were determined. MIC50 and MIC90 were determined by identifying the MIC which inhibits 50% or 90% of the collection. Clinical breakpoints from EUCAST (2021) were used where applicable and cut off points for CHX and OCT were set at 4 μg/mL and 2 μg/mL respectively, as defined by Htun et al. (2019). Statistical analysis was performed using Graph Pad Prism (v5.04). For analysis of more than 2 groups a 1-way ANOVA Kruskal-Wallis, Non-Parametric test was used.

### Sensitivity to pH

Two randomly selected isolates from each of the Non NRCS-A, proto NRCS-A and NRCS-A groups were used to test for ability to grow at pH (pH range 3.5 −10). Overnight cultures of each were used to inoculate plates in duplicate (PM10 MicroPlate™ pH, Biolog, U.K) in accordance with the manufacturer’s protocol and respiratory activity in each sample measured over 24 hours automatically using the OmniLog reader (Biolog, U.K).

## Results

### *Staphylococcus capitis* were isolated from various body sites of babies on NICUs

Routine MRSA swabs were taken from a total of 159 babies across their stay in NICU, *S. capitis* was isolated from 55 babies (34.5 %) admitted to NICU in the UK and Germany yielding a total of 102 *S. capitis* isolates (UK n=90, German n=14) from 4 different body sites (ear n=47; gut n=13; nose n=17; skin n=27). A further 12 isolates were from neonates with infection, 11 from blood and one from skin. A total of 15 *S. capitis* were collected from adults including from prosthetic joint infection (n=3), blood (n=5) healthy skin swabs (n=6) and one from an abdominal swab. All these 129 strains were sequenced and gave an average genome size of 2.52 Mbp with an average GC content of 32.86 %. A further 9 previously published reference genomes were also included in later analysis of genomic relationships.

### Antiseptic susceptibility differed between isolates from different geographical sites

All the isolates from the NICU CoNS collection were tested for antiseptic and antibiotic susceptibility (Supplementary Data 2, Sethi et al. (2021)). The *S. capitis* isolates demonstrated greater sensitivity to octenidine than chlorhexidine (MIC^50^ of 2 μg/mL and 16 μg/mL respectively, and MIC^90^ being 4 μg/mL and 32 μg/mL) (Figure 1). There was no significant difference found in susceptibility of isolates from different body sites, but the MIC^50^ values for both octenidine and chlorhexidine in the German isolates (OCT 1 μg/mL; CHX 8 μg/mL) were lower than those of the UK isolates (OCT 2 μg/mL; CHX 16 μg/mL). As well as the antiseptics, UK isolates had higher MIC^50^ values for gentamicin, penicillin and fusidic acid. The largest discrepancy being for fusidic acid where the MIC^50^ increased from 0.625 μg/mL for the German population to 4 μg/mL in UK isolates. Seven adult isolates showed hetero-resistance to daptomycin (defined as an MIC ≥ 1 μg/mL, Becker et al. (2014)) but this was not seen in any neonatal isolates. No vancomycin resistance was observed in the isolates with the highest MIC being 2 μg/mL, however nearly a quarter of the isolates exhibited heteroresistance to vancomycin (29 in total), with the majority of these (23/29) being from NICU.

**Figure 1.**
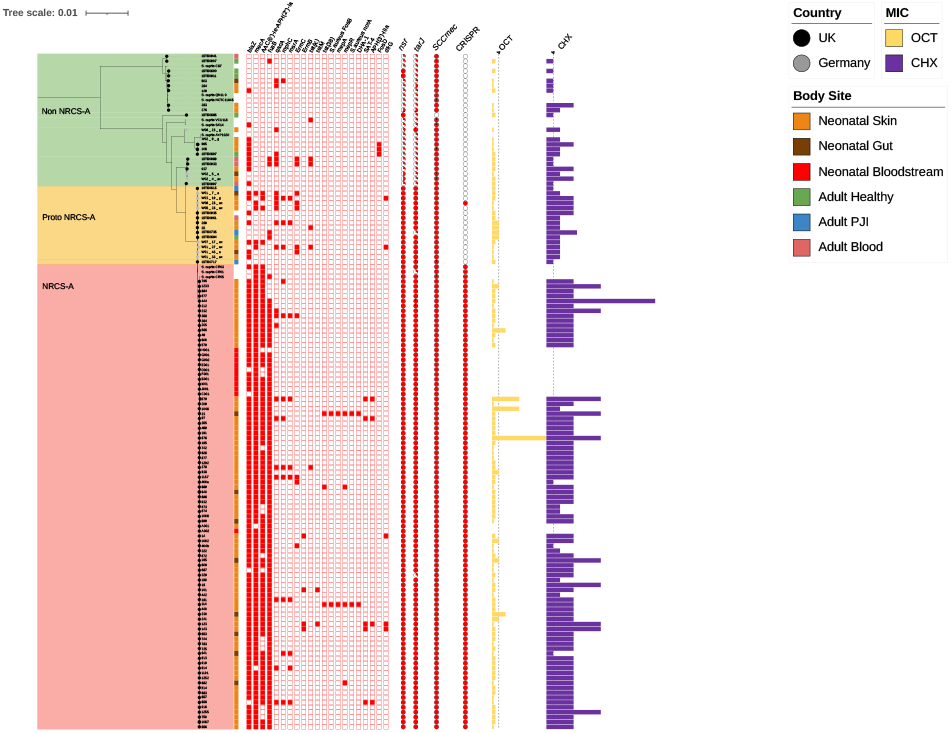
Population structure of *S. capitis*. A maximum likelihood (ML) tree based on core gene alignment from Roary. Three groups are indicated; Non NRCS-A isolates are highlighted in green, proto NRCS-A in orange, and NRCS-A, in pink. Country of isolation is indicated by the colour of tip branches; Germany (grey circles), UK (black circles). Body site of isolation are shown by coloured boxes; orange – neonatal skin, brown – neontal gut, red – neonatal bloodstream, green – healthy adult skin, blue – adult prosthetic joint, pink – adult bloodsteam. Antimicrobial resistance (AMR) genes are represented by red boxes (blank boxes represent absence of the corresponding gene). The percentage identity of *nsr* and *tarJ* in each strain (compared to the reference sequences present in CR01 and CR05 respectively using BLASTp) are shown as pie charts. MIC data for chlorhexidine (CHX, purple bars) and octenidine (OCT, yellow bars) are shown with the dotted lines representing 4 μg/mL of OCT or CHX.

### Phylogenetic characterisation of *S. capitis* identified two major clades

A phylogenetic tree was built using the core gene alignment output by Roary of all isolates in the panel (as well as the nine available NCBI reference *S. capitis* genomes in Supplementary Data 1) then aligned with IQTREE (Figure 1). Clades including NRCS-A strains were determined based on similarity to known NRCS-A reference strains (CR01, 03 and 05) and by the presence of the two genes *nsr* and *tarJ*, common identifiers used for this clone (Martins Simões et al., 2013). The tree showed a diversity of non-NRCS-A clones and then a clonal group which all carried *nsr* and *tarJ* and included the NRCS-A strains but could be divided into two further groups based on carriage of a CRISPR system.

**The ‘non NRCS-A’** group (Figure 1, green, n=27) contained no known NRCS-A strains but did include other *S. capitis* NCBI reference genomes and isolates from adults as well as some NICU isolates. Isolates in this group tended to show a low tolerance to OCT and CHX (OCT MIC_50_ = 2 μg/mL, CHX MIC_50_ = 8 μg/mL; Table 1 and fusidic acid with *fusB* being absent in all but one strain (Supplementary Data 3). Of these non NRCS-A isolates 12 came from the NICU (4 German, 8 UK), nine came from healthy adult skin swabs (n=5) or blood cultures (n=4) and this group also included six non NRCS-A reference strains (*S. capitis* strains AYP1020, C87, QN1, SK14, VCU116 and NCTC11045 (Supplementary Data 1)).

**Table 1.** MIC^50^- and MIC^90^ values of antimicrobials against the *S. capitis* groups.

|  | OCT |  | CHX |  | VAN |  | GEN |  | PEN |  | CEF |  | CIP |  | DAP |  | FUS |  |
| --- | --- | --- | --- | --- | --- | --- | --- | --- | --- | --- | --- | --- | --- | --- | --- | --- | --- | --- |
|  | MIC<sup>50</sup> | MIC<sup>90</sup> | MIC<sup>50</sup> | MIC<sup>90</sup> | MIC<sup>50</sup> | MIC<sup>90</sup> | MIC<sup>50</sup> | MIC<sup>90</sup> | MIC<sup>50</sup> | MIC<sup>90</sup> | MIC<sup>50</sup> | MIC<sup>90</sup> | MIC<sup>50</sup> | MIC<sup>90</sup> | MIC<sup>50</sup> | MIC<sup>90</sup> | MIC<sup>50</sup> | MIC<sup>90</sup> |
| Non NRCS-A | 2 | 2 | 8 | 16 | 1 | 2 | 0.064 | 32 | 0.5 | 8 | 1 | 4 | 0.5 | 4 | 0.5 | 2 | 2 | 2 |
| Proto NRCS-A | 2 | 4 | 8 | 16 | 1 | 2 | 1 | 32 | 2 | 32 | 1 | 4 | 0.25 | 0.5 | 0.5 | 1 | 0.064 | 2 |
| NRCS-A | 2 | 4 | 16 | 32 | 1 | 2 | 16 | 32 | 8 | 126 | 4 | 4 | 0.5 | 4 | 0.25 | 1 | 4 | 4 |
| Overall | 2 | 4 | 16 | 16 | 1 | 2 | 8 | 32 | 4 | 126 | 4 | 4 | 0.5 | 4 | 0.5 | 1 | 2 | 4 |
| UK | 2 | 4 | 16 | 32 | 1 | 2 | 8 | 32 | 4 | 126 | 4 | 4 | 0.5 | 4 | 0.5 | 1 | 4 | 4 |
| German | 1 | 2 | 8 | 16 | 1 | 1 | 8 | 32 | 4 | 126 | 4 | 4 | 0.5 | 4 | 0.5 | 1 | 2 | 2 |

**The ‘Proto NRCS-A’** group (denoted as such as it appears ancestrally related to the NRCS-A group and contain *nsr* and *tarJ* but no CRISPR system) (Figure 1, orange, n=16) carried fewer AMR genes than the NRCS-A isolates and typically lacked *fusB*, however these strains were closely related to the NRCS-A group. This proto-NRCS-A clade had OCT and CHX susceptibilities lower than NRCS-A (OCT MIC50 = 2 μg/mL, CHX MIC50 = 8 μg/mL: Table 1). Along with ten isolates from NICU, the remaining isolates were sourced from adult PJI (n=3), blood (n=1), an abdominal swab and a healthy adult skin swab.

**The ‘NRCS-A’** group (Figure 1, red, n=95) was the largest and contained most isolates as well as NRCS-A reference strains, *S. capitis* CR01, CR03 and CR05 along with the remaining *S. capitis* isolated from the NICU. No isolates from adults were found in this clade. Tolerance to OCT and CHX was marginally higher than isolates from proto NRCS-A (OCT MIC_50_ = 2 μg/mL, CHX MIC_50_ = 16 μg/mL: Table 1). NRCS-A was defined by isolates carrying multiple AMR genes including *blaZ* (β-lactamase)*, fusB* (fusidic acid resistance)*, AAC(6’)-la-APH(2”)-la* (aminoglycoside resistance) and *mecA* (penicillin/methicillin resistance) and contained the *nsr* and *tarJ* genes as well as a CRISPR-Cas Type III-A system (Figure 1 and Supplementary Data 3).Multiple isolates of *S. capitis* were isolated on different sampling dates from 24 separate babies (Figure 2) with the remaining 40 babies only yielding a single isolate. Only 4/64 babies carried only non NRCS-A *S. capitis* strains (babies 48, 58, 94 and 146, Figure 2). The remaining 60 babies carried NRCS-A isolates (Supplementary Data 2). Three babies (baby 8, 59, and 137) yielded *S. capitis* isolates from two of the three groups (Non NRCS-A 1 and NRCS-A), and one baby had two distinct isolates from the proto NRCS-A group (baby 28), and another yielded one non NRCS-A and one proto NRCS-A isolate (Supplementary Data 2).

**Figure 2.**
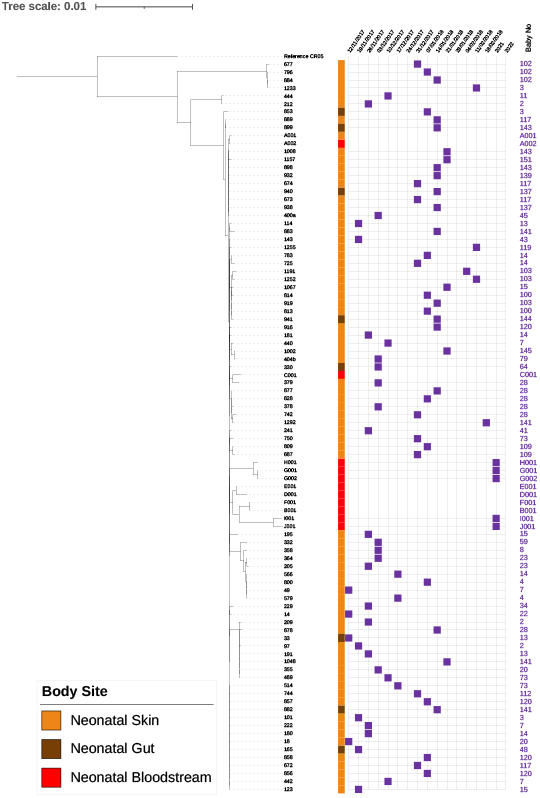
Detailed phylogeny of NRCS-A based on core SNP alignment. The week of isolation is indicated by purple squares for all UK isolates and sources of each isolate are indicated by coloured squares (neonatal skin = orange; neonatal gut = brown, neonatal bloodstream = red).

The number of SNPs distinguishing all 138 isolates was determined using snippy with UK NRCS-A CR05 as a reference. The mean (standard deviation) number of SNPs seen between isolates in the NRCS-A, and proto NRCS-A groups were 115.47 and 265 respectively (Supplementary Data 4). On average the non NRCS-A isolates contained 23084.5 (3263.2) SNPs, a much higher number confirming that this clade is genetically distant from the NRCS-A isolates and more diverse. The 92 NRCS-A isolates from NICU had a maximum of 31 snps between each other (mean aligned core genome: 2401507 bp), these isolates were only isolates from the UK (Supplementary Data 4). This high degree of relatedness of isolates in the UK NICUs suggests a recent common ancestor with transition across the unit and further confirms the nature of NRCS-A.

### Evidence for transmission within the NICU

We analysed relationships between isolates taken longitudinally from babies (Figure 2), this revealed a cluster of 8 isolates (209, 878, 33, 97, 191, 1048, 355, 489) differentiated by only 3 SNPS, (Supplementary Data 5) recovered between over 70 days, from six different babies (Figure 2) suggesting transmission within the NICU environment and persistence of isolates over time. The total number of SNPs between all NRCS-A isolates was only 37. This indicates that very similar closely related isolates were present in the UK NICUs and they remain present for years with very little genomic change.

### Differences distinguishing clades

The collection was analysed by Roary, (Page et al., 2015) to identify genes that were either present or absent in the clades, ‘Scoary’ was then used to calculate statistical associations between the presence and absence of genes, clade membership and a series of phenotypes (Brynildsrud et al., 2016). Notably the genes being most significantly associated with the NRCS-A clade were the CRISPR genes, all NRCS-A isolates harboured a complete CRISPR-Cas Type-III-A system. None of the closely related proto NRCS-A isolates harboured any CRISPR Cas Type-III-A genes. These Cas Type-III-A genes are inserted in the *SCCmec-SCCcad/ars/cop* element (Figure 3). Two virulence genes *paaZ* and *ugpQ*, along with *dut* are known to be closely located to *mecA*, and were also strongly associated with being present in the NRCS-A isolates (Datta et al., 2021, Westberg et al., 2022).

**Figure 3.**
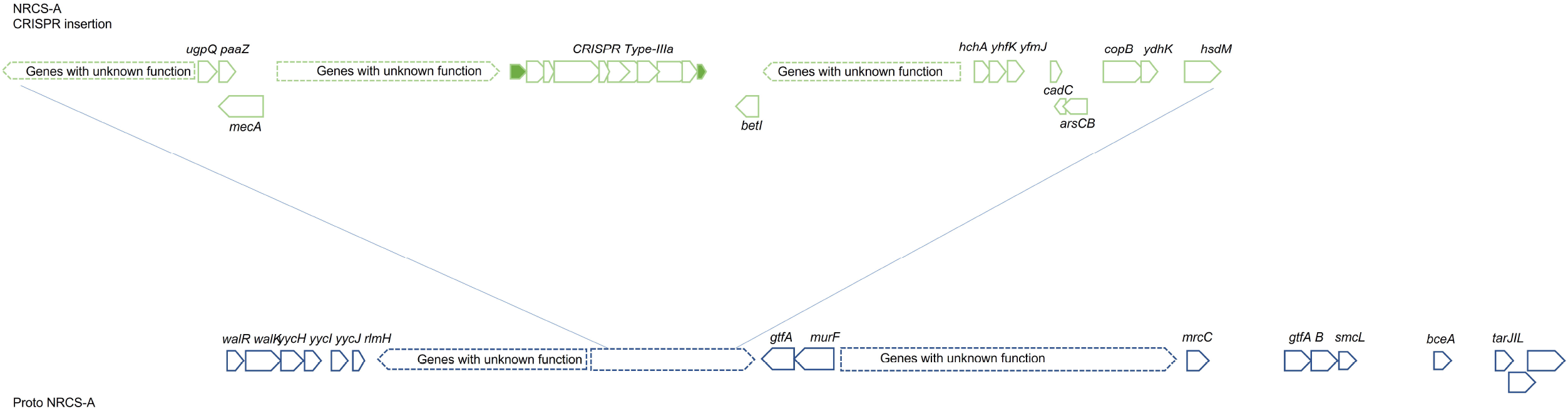
Insertion of the CRISPR-Cas Type III-A system in NRCS-A isolates, compared to the related Proto NRCS-A isolates.

Scoary analysis also identified groups of other genes significantly associated with the NRCS-A isolates. Many of these genes were annotated as being of unknown function and clustered together in the genomes. Further analysis of these genes, using Blastn identified several as being putative phage terminases which are inserted in-between *yfnK* and *murI*, immediately downstream from *frdA, uvrC* and *mutS*. Scoary also identified *mcrC* in NRCS-A isolates which is involved in regulation of degrading methylated bacteriophage DNA (Nirwan et al., 2019) (Table 2, Supplementary Data 6).

**Table 2.**
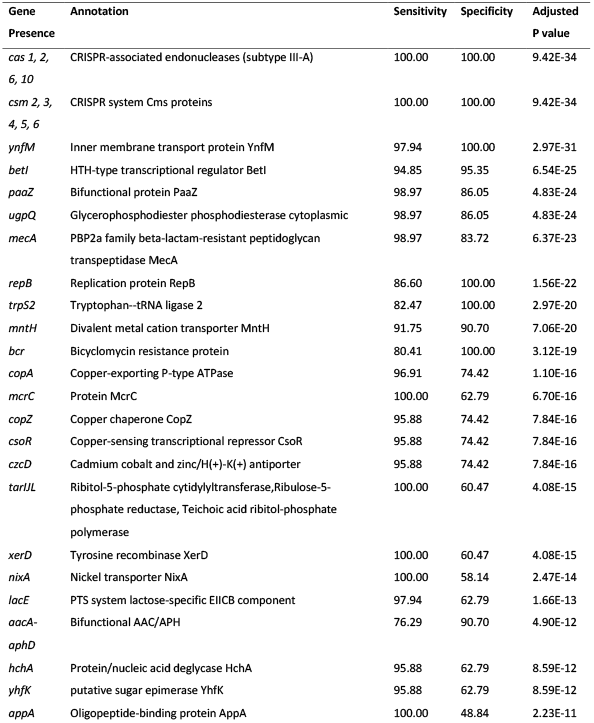
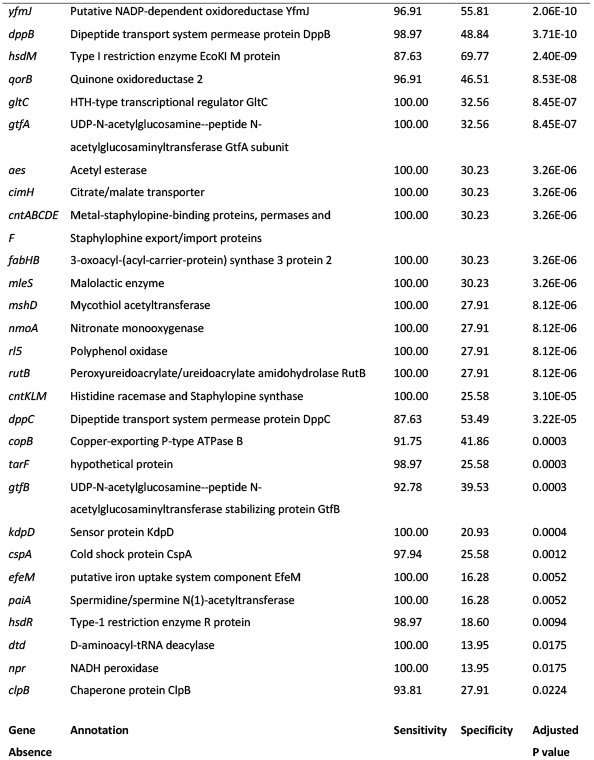
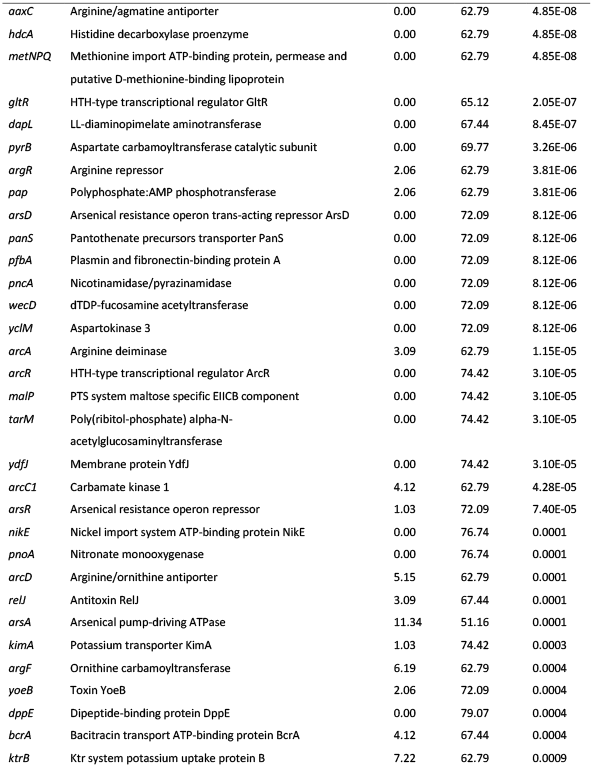
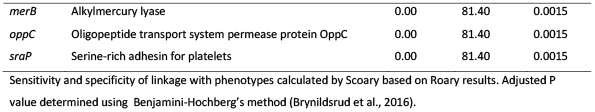
Selected genes where presence/absence was strongly statistically associated with the NRCS-A isolates compared other *S. capitis* clades.

Previous research by Martins Simões et al. (2013) suggested the presence of *nsr* and *tarJ* as a defining characteristic of the NRCS-A clone, and both these genes are present in both NRCS-A and proto NRCS-A isolates. The *tarLIJ* locus encodes a Ribulose-5-phosphate reductase which are involved in teichoic acid biosynthesis (Tevell et al., 2020, Martins Simões et al., 2013). These were significantly associated with the NRCS-A group, parts of the operon were found in Non NRCS-A but were usually incomplete (Figure 1, Table 2). All NRCS-A and proto NRCS-A isolates had *tarJ* compared to only 3.7% of non NRCS-A isolates (Figure 1). The *nsr* gene associated with nisin resistance was identified in both NRCS-A and proto NRCS-A isolates, and *kdpDC* was also significantly over-represented in these groups which encodes a potassium pump encoded close to *nsr* in the genome (Martins Simões et al., 2013).

There was a significant increase in the presence of genes involved in metal transport in the NRCS-A group compared to the others (Table 3, Supplementary Data 6). ABC permeases *cnt ABCDEFKL, nikC* and *oppDF* were all identified and are involved in the acquisition of Zn and Ni (Grim et al., 2017). Further genes involved in metal detoxification were also seen in NRCS-A, the *copB, copAZ, arsBC* and *cadC*systems were over-represented in this clade. These have all been previously identified as part of an *SCCmec* element (*SCCmec-SCCcad/ars/cop*) also containing the CRISPR-Cas Type III-A system in NCRS-A isolates. As well as genes associated with this element, the NRCS-A isolates could also be distinguished from the proto-NRCS-A clade by the presence of *mntH* (Table 3, Supplementary Data 6).

**Table 3.** Selected genes where presence was strongly statistically associated with the NRCS-A isolates compared to the proto-NRCS-A isolates.

| Gene | Annotation | Sensitivity | Specificity | P value |
| --- | --- | --- | --- | --- |
| *cas1, 2, 6, 10* | CRISPR-associated endonucleases (type III-A) | 100.00 | 100.00 | 1.81E-16 |
| *csm2,3,4,5,6* | CRISPR system Cms proteins | 100.00 | 100.00 | 1.81E-16 |
| *bceA* | Bacitracin export ATP-binding protein BceA | 100.00 | 93.75 | 6.26E-15 |
| *ynfM* | Inner membrane transport protein YnfM | 97.94 | 100.00 | 6.26E-15 |
| *smc* | Chromosome partition protein Smc | 96.91 | 100.00 | 3.07E-14 |
| *npr* | NADH peroxidase | 95.88 | 100.00 | 1.27E-13 |
| *lytO* | putative autolysin LytO | 95.88 | 100.00 | 1.27E-13 |
| *xerD* | Tyrosine recombinase XerD | 96.91 | 93.75 | 1.60E-12 |
| *araB* | Ribulokinase | 92.78 | 100.00 | 4.57E-12 |
| *cmoM* | tRNA 5-carboxymethoxyuridine<br>methyltransferase | 90.72 | 100.00 | 3.45E-11 |
| *lexA* | LexA repressor | 90.72 | 100.00 | 3.45E-11 |
| *dut* | Deoxyuridine 5'-triphosphate<br>nucleotidohydrolase | 88.66 | 100.00 | 1.87E-10 |
| *betI* | HTH-type transcriptional regulator BetI | 94.85 | 87.50 | 7.21E-10 |
| *repB* | Replication protein RepB | 86.60 | 100.00 | 9.40E-10 |
| *trpS2* | Tryptophan--tRNA ligase 2 | 82.47 | 100.00 | 1.52E-08 |
| *mntH* | Divalent metal cation transporter MntH | 91.75 | 87.50 | 1.71E-08 |
| *hisC* | Histidinol-phosphate aminotransferase | 81.44 | 100.00 | 2.71E-08 |
| *bcr* | Bicyclomycin resistance protein | 80.41 | 100.00 | 4.93E-08 |
| *blaR1-2* | beta-lactam sensor/signal transducer BlaR1 | 95.88 | 75.00 | 7.08E-08 |
| *hin* | DNA-invertase hin | 95.88 | 75.00 | 7.08E-08 |
| *ldh1* | L-lactate dehydrogenase 1 | 78.35 | 100.00 | 1.41E-07 |
| *mecA* | PBP2a family beta-lactam-resistant<br>peptidoglycan transpeptidase MecA | 98.97 | 62.50 | 1.59E-07 |
| *paaZ* | Bifunctional protein PaaZ | 98.97 | 62.50 | 1.59E-07 |
| *ugpQ* | Glycerophosphodiester phosphodiesterase<br>cytoplasmic | 98.97 | 62.50 | 1.59E-07 |
| *blal* | Penicillinase repressor | 93.81 | 75.00 | 6.17E-07 |
| *rhaS* | HTH-type transcriptional activator RhaS | 79.38 | 93.75 | 2.38E-06 |
| *blaZ* | Beta-lactamase | 84.54 | 87.50 | 2.55E-06 |
| *copA* | Copper-exporting P-type ATPase | 96.91 | 62.50 | 3.14E-06 |
| *copZ* | Copper chaperone CopZ | 95.88 | 62.50 | 9.94E-06 |
| *dtd* | D-aminoacyl-tRNA deacylase | 83.51 | 81.25 | 5.74E-05 |
| *menH* | 2-succinyl-6-hydroxy-2 4-cyclohexadiene-1-carboxylate synthase | 93.81 | 62.50 | 6.91E-05 |
| *ydhP* | Inner membrane transport protein YdhP | 90.72 | 68.75 | 7.62E-05 |
| *csoR* | Copper-sensing transcriptional repressor CsoR | 95.88 | 56.25 | 9.74E-05 |
| *czcD* | Cadmium cobalt and zinc/H(+)-K(+) antiporter | 95.88 | 56.25 | 9.74E-05 |

Sensitivity and specificity of linkage with phenotypes calculated by scoary based on Roary results. Adjusted P value (<0.001) determined using Benjamini-Hochberg's method (Brynildsrud et al., 2016).

Previously it has been suggested that NRCS-A isolates are adapted to survive in both the gut and on the skin, two environments where pH differs. We observed that growth of both proto NRCS-A and NRCS-A isolates at lower pH was higher compared to non NRCS-A isolates (Figure 4). Growth at pH 6 and pH 5.5 was also marginally higher compared to pH 7 in both the NRCS-A and proto NRCS-A isolates (Figure 4).

**Figure 4.**
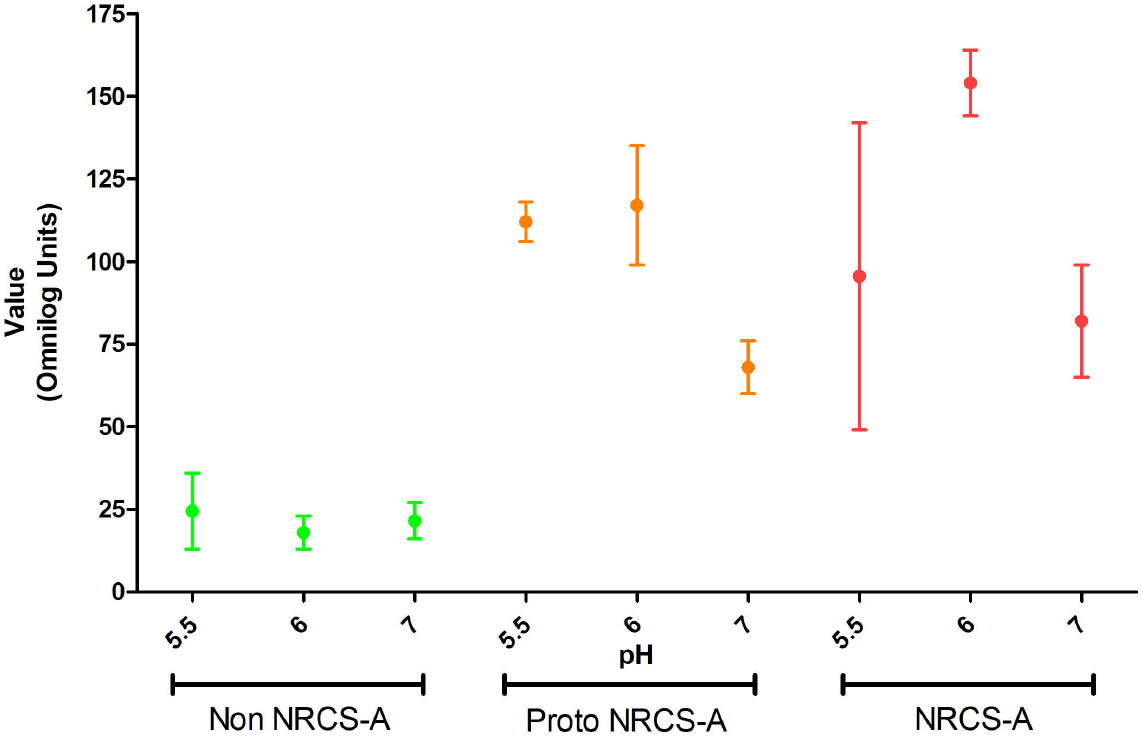
Growth of *S. capitis* in different pH (pH7, 6 and 5.5), showing growth at final time point of 48 hours from PM10 MicroPlate ^™^. Points indicate mean values and bars the standard error.

To compare conservation of the *SCCmec* element seen in all the isolates, the *SCCmec-SCC-cad/ars/cop* sequence of CR01 (GenBank KF049201.1) was compared to all the *S. capitis* isolates using snippy-core and snp-dist (Seemann, 2015, Seeman, 2019). All isolates and references except for VCUII6 aligned to the *SCCmec-SCC-cad/ars/cop* reference. The NRCS-A isolates, on average, had 12.3 ± 0.5 SNPs, which increased to 64.4 ± 11.1 in non NRCS-A and 190.9 ± 22.5 in proto NRCS-A isolates. The higher divergence in the proto NRCS-A group is surprising given its closer relationship to the NRCS-A group according to the core genome phylogeny.

Further analysis of the CRISPR-Cas Type III-A system showed that the genes were highly conserved when aligned. In all isolates it was possible to identify direct repeat (DR) sequences (beginning with GATAACT), and there were three different DRs present in the isolates as previously observed in CR01 Cao et al. (2016). In a full SNP ClustalW alignment DR5 (4 times) and DR6 (3 times) were found before the Cas genes, and then two repeats of DR8 towards the end of the alignment after the CRISPR genes (Cao et al., 2016).

### Carriage of AMR genes varied between non NRCS-A and NRCS-A

All isolates contained *lmrS, norA, arlR* and *dfrC*, and all but one isolate contained *mgrA*. NRCS-A was defined by the additional presence of *blaZ, mecA, ACC(6’)-le(APH)(2’)-la* and *fusB* in the majority of isolates whereas proto NRCS-A had a smaller number of AMR genes which were interspersed across the clade without a consistent pattern (Figure 1, Supplementary Data 3). Analysis of the pangenomes with Scoary was used to identify genes correlating with susceptibility to different antimicrobials. Unsurprisingly *mecA* was strongly associated with high cefotaxime MICs (4 μg/mL, Supplementary Data 6). Carriage of *blaZ* was also associated with penicillin resistance (≥1 μg/mL, P= 0.005), however CRISPR genes (P= 0.002) and other genes commonly found on the *SCCmec-SCC-cad/ars/cop* MGE were more significant (Table 4). The *qoc* family of genes has previously been implicated in tolerance to antiseptics, *qacA* was detected but only present in 22/138 (16 %) isolates and there was no correlation with the presence of *qacA* and CHX tolerance. Carriage of *qacA* was only observed in 16.25% of isolates with high chlorhexidine MICs ≥16 μg/mL (Figure 1). For isolates that had an MIC to chlorhexidine of ≥16 μg/mL the genes mostly associated with this tolerance were the CRISPR-Cas Type III-A, Supplementary Data 6). This is a marker of NRCS-A isolates which had significantly higher chlorhexidine MICs (Figure 5) although there was no obvious genetic basis for this. Scoary was unable to identify any significant genes associated with octenidine susceptibility.

**Table 4.** Selected genes where presence was statistically associated with antibiotics and chlorhexidine susceptibility.

| Gene | Annotation | Adjusted P value |  |  |
| --- | --- | --- | --- | --- |
|  |  | CHX<br>16 µg/mL | PEN<br>1 µg/mL | CEF<br>4 µg/mL |
| *aacA-aphD* | Bifunctional AAC/APH | 0.0001 |  |  |
| *araB* | Ribulokinase | 2.47E-06 |  |  |
| *bcr* | Bicyclomycin resistance protein | 7.49E-06 |  |  |
| *betI* | HTH-type transcriptional regulator BetI | 1.57E-06 |  |  |
| *bin3* | Putative transposon Tn552 DNA-invertase bin3 | 2.79E-06 | 0.002 | 0.0126 |
| *blaI* | Penicillinase repressor | 3.50E-05 |  |  |
| *blaR1-2* | beta-lactam sensor/signal transducer BlaR1 | 0.000414331 |  |  |
| *cas1,2,6,10* | CRISPR-associated endonuclease Cas1 | 1.25E-07 | 0.002 | 0.0126 |
| *cls* | Cardiolipin synthase | 7.97E-05 |  |  |
| *copA* | Copper-exporting P-type ATPase | 3.44E-06 | 0.002 | 0.0153 |
| *copZ* | Copper chaperone CopZ | 3.44E-06 | 0.002 | 0.0153 |
| *csm2,3,4,5,6* | CRISPR system Cms protein Csm2 | 1.25E-07 | 0.002 | 0.0126 |
| *csor* | Copper-sensing transcriptional repressor CsoR | 3.44E-06 |  |  |
| *czcD* | Cadmium cobalt and zinc/H(+)-K(+) antiporter | 1.35E-06 | 0.003 |  |
| *dut* | Deoxyuridine 5'-triphosphate nucleotidohydrolase | 7.83E-07 | 0.002 |  |
| *hin* | DNA-invertase hin | 2.79E-06 | 0.002 | 0.0126 |
| *hisC* | Histidinol-phosphate aminotransferase | 7.49E-06 |  |  |
| *ldh1* | L-lactate dehydrogenase 1 | 3.06E-05 |  |  |
| *lexA* | LexA repressor | 2.47E-06 | 0.002 |  |
| *lytO* | putative autolysin LytO | 4.90E-05 |  |  |
| *mbolR* | hypothetical protein | 1.25E-07 |  |  |
| *mecA* | PBP2a family beta-lactam-resistant peptidoglycan<br>transpeptidase | 1.20E-06 |  | 0.0126 |
| *mntH* | Divalent metal cation transporter MntH | 2.53E-06 |  | 0.0126 |
| *npr* | NADH peroxidase | 5.22E-06 |  |  |
| *paaZ* | Bifunctional protein PaaZ | 3.34E-07 |  | 0.0126 |
| *repB* | Replication protein RepB | 0.0001 |  |  |
| *rhaS* | HTH-type transcriptional activator RhaS | 1.02E-05 |  |  |
| *rihC* | Non-specific ribonucleoside hydrolase RihC |  | 0.005 |  |
| *smc* | Chromosome partition protein Smc | 1.25E-07 | 0.002 | 0.0126 |
| *trpS2* | Tryptophan--tRNA ligase 2 | 2.79E-06 |  |  |
| *ugpQ* | Glycerophosphodiester phosphodiesterase<br>cytoplasmic | 3.34E-07 |  | 0.0126 |
| *xerC* | Tyrosine recombinase XerC | 1.25E-07 | 0.002 |  |
| *xerD* | Tyrosine recombinase XerD | 1.24E-06 |  |  |
| *ynfM* | Inner membrane transport protein YnfM | 1.25E-07 | 0.002 | 0.0126 |

**Figure 5.**
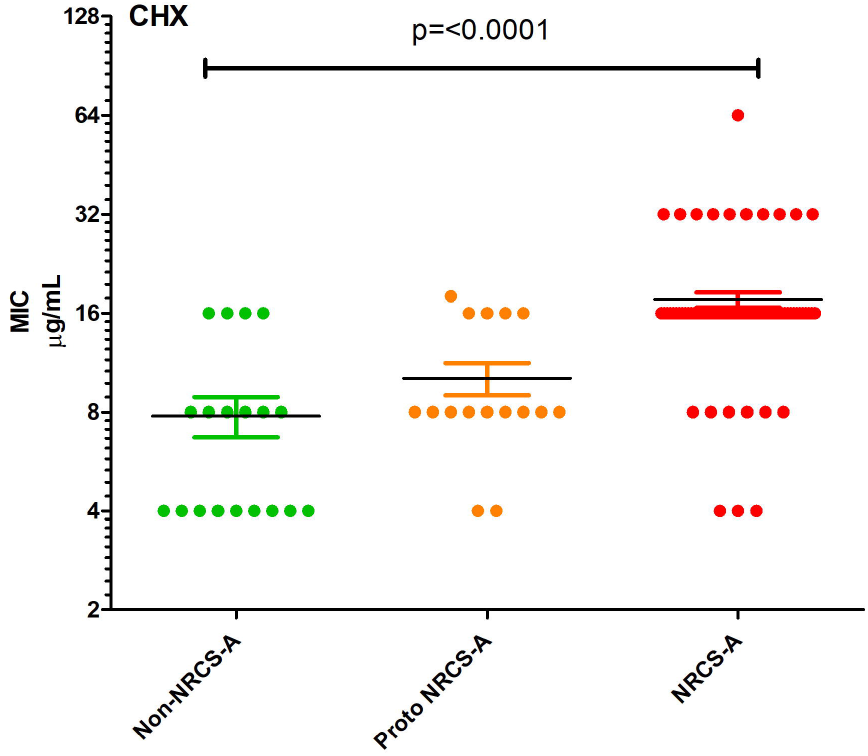
Chlorhexidine susceptibility of the three groups of *S. capitis*. Points represent values from individual strains, horizontal lines indicate the means with bars showing the standard error of the mean.

The classic antibiogram for NRCS-A isolates includes being resistant to fusidic acid (PHE, 2021), with the clinical breakpoint being ≥0.5 μg/mL for all CoNS (EUCAST, 2021). Based on this criterion 79% of our collection tested were clinically resistant and carriage of *fusB* was seen in 78.75 % of these resistant isolates, leaving 17 isolates with no known mechanism of fusidic acid resistance. The presence of *fusB* was mainly observed in NRCS-A, where there was a small increase in tolerance to fusidic acid, with the mean MIC being 2.95 (SD: 0.155) μg/mL, higher than non NRCS-A (1.40, SD: 0.301 μg/mL). The UK isolates had a higher median FUS MIC (4 μg/mL) than the German isolates (1.5 μg/mL) (P = <0.01). Penicillin resistance encoded by *mecA* was seen in 86% of the isolates with an MIC ≥ 2 μg/mL (Supplementary Data 2 and 3), and carriage of *ACC(6’)-la-APH(2”)-la* was seen in 73% of isolates with an MIC ≥1 μg/mL of gentamicin. No genes were associated with ciprofloxacin resistance although the main mechanism of resistance to this agent is target site mutation within *gyrA* rather than the presence of a gene.

*S. capitis* NRCS-A are often referred to as demonstrating hetero-resistance to vancomycin, indicated by a MIC of 2 μg/mL (Satola et al., 2011). Despite 23.5% of the isolates having an MIC of 2 μg/mL there was no presence of *vanA* apart from in one isolate (with an MIC of 1 μg/mL) or any other significant genes associated with vancomycin resistance.

## Discussion

Even with the increased focus on their prevention, nosocomial infections remain a major cause of disease and death in hospitals. Transmission of bacterial pathogens can occur via multiple routes and vectors with both health care workers and the environment sources of infection for many pathogens such as *Staphylococcus aureus, Pseudomonas aeruginosa* and Enterobacteriaceae (Risso et al., 2019, Götting et al., 2020). The effective adaption of the NRCS-A clone to the NICU environment is well established, though the reasons for this remain unclear. One suggestion is that NRCS-A isolates are adapted to survival in the gut which provides an additional reservoir to the skin and may allow persistent colonisation, even when skin organisms are lost during antisepsis (Butin et al., 2017). Carriage of the *nsr* gene responsible for nisin resistance has been proposed to aid survival in the gut where this antimicrobial peptide is produced (Martins Simões et al., 2013). In this study we documented carriage of NRCS-A on both the skin of neonates and in the gut, and this was much more common than for other groups of *S. capitis* (Figure 1). We did observe that the *nsr* gene is present in NRCS-A isolates but not in more distantly related strains. Interestingly we identified a group of isolates closely related to NCRS-A, the ‘proto NRCS-A’ strains which did carry *nsr* but which were isolated from the gut less often (Figure 1). It was also observed that the proto NRCS-A and NRCS-A isolates, which possess *nsr* and *tarJ*, had increased growth in pH 6-5.5 cultures (Figure 4), analogous to their niche in the gastrointestinal tract.

### Metal acquisition and survival

Analysis of the pangenomes of each group identified other genes that may aid the survival of the NRCS-A in the NICU. In the NRCS-A isolates there was a marked increase in genes associated with metal chelation (Table 2), availability of many metals is limited *in vivo* and in the gut metals are precious commodities. Metal chelators can also influence virulence, for example, the nickel/cobalt transporter *nixA* in *S. aureus* has been implicated in an increase in virulence in UTI and kidney infection and is required for urease activity in mice (Remy et al., 2013). Other metal related genes found in NRCS-A were *mntH*, a metal transporter found to minimise impact of Mn and Zn starvation (Kehl-Fie et al., 2013), and *cntABCDEF* which functions with Staphylopine, StP (CntKLM) a staphylococcal zincophore involved in Zn acquisition which can also bind to Cu, Co and Ni (Grim et al., 2017, Song et al., 2018). These high affinity chelators and zincophores have been shown to be able to outcompete human calprotectin (a Zn chelator excreted by neutrophils) to evade nutritional immunity. The presence of *cntABCDEF cntKLM, nixA* and *mntH* may indicate that even in nutritionally deprived environments, the NRCS-A isolates are able to acquire essential metal ions and survive in a gut environment. Similarly, the presence of *copA, cad*, and *ars*, encoded within the *SCCmec-SCCcad/ars/cop* element in NRCS-A provide defence against toxic metals by encoding efflux systems. Together these data suggest that an enhanced ability to scavenge and survive metal exposure may be crucial for the success of NRCS-A and supports the theory that gut colonisation is a key strategy exploited by this clone.

### Phage defence

If NRCS-A has developed a lifestyle which includes ability to colonise the gut then it also needs to adapt to attack from phages, which are prevalent in this environment. In *Escherichia coli* the *mcrBC* genes provide protection from phages and McrC activates McrB to degrade methylated bacteriophage DNA (Nirwan et al., 2019). MrcC was seen in our collection but *mcrB* was not detected and it is unclear what impact *mcrC* alone may have. Other mechanisms of phage defence observed in the NRCS-A isolates were the CRISPR-associated genes (Cas), used in defence against MGEs, viruses and plasmids (Tamulaitis et al., 2017, Rossi et al., 2017). CRISPR-Cas systems show low abundance in most Non-aureus Staphylococci although they have been documented in the genomes of *S. aureus* strains as well as in *S. epidermidis, S. schleiferi, S. haemolyticus, S. lugdunensis*. Carriage of a CRISPR-Cas system is a known feature of NRCS-A *S capitis* and CRISPR-Type III-A Cas genes were identified in all NRCS-A isolates. This was the main defining feature differentiating the proto NRCS-A and the NRCS-A clades, however when this system was acquired by NCRS-A remains unclear. The direct repeats identified are similar to that as previously seen in CR01 and CR03 which both contain the conserved CCCC and GGGG motifs which are thought to form hairpin structures (Rossi et al., 2017). The highly conserved regions suggest that a common ancestor is responsible for the acquisition of the CRISPR-Cas Type-III-A genes.

### Antiseptic tolerance

One defining feature of the NRCS-A clade was increased tolerance of chlorhexidine (Figure 5), this was not associated with carriage of *qac* genes, and the mechanism remains unclear. This increased ability to survive chlorhexidine exposure may allow both survival of skin antisepsis but also environmental decontamination where chlorhexidine is a common part of cleaning and disinfection routines for incubators and medical equipment. There was no elevation in tolerance to octenidine, which is the main alternative for antisepsis in neonates (Sethi et al., 2021).

### Stability of NRCS-A

The NRCS-A isolates compared here were very similar including those separated by geography (UK and German isolates) and time (reference genomes spanning years in isolation) (Figure 2), and there were no more than 37 SNPs separating strains in the core genome. We provide evidence for transfer between babies on the NICU (Figure 2), studies identifying NRCS-A in the general environment and on different surfaces or medical implements following clusters of LOS NRCS-A positive cases have suggested a role for the environment in transmission The isolates from carriage and the 10 from neonatal blood cultures were highly similar (Figure 1). This supports the idea that carriage acts as a reservoir for infection but clearly carriage by neonates in a nosocomial setting does not usually result in infection.

## Conclusions

We have identified that carriage of the NRCS-A clone is abundant on the skin and gut of uninfected neonates and that they are likely to be transferred within NICUs. NRCS-A isolates separated in time and space showed little genetic variation and carriage isolates were indistinguishable to those from blood culture, suggesting carriage can be a precursor to infection. We identified a closely related proto-NRCS-A clade which was less associated with gut carriage, did not possess the CRISPR-Cas III-A system, demonstrated lower antimicrobial resistance and chlorhexidine tolerance and carried fewer metal acquisition or detoxification genes. Our data supports the idea that gut colonisation is a key survival strategy of NRCS-A and expands our understanding of the likely mechanisms employed by this clone to both survive on skin, in the environment and in the gut. This opens the possibility to develop a probiotic approach to hinder colonisation of NCRS-A based upon nutritional immunity where strains that sequester essential metals may prove beneficial to the infant gut. Recent deployment of a probiotic product into the NNUH NICU has been used to reduce incidence of necrotising enterocolitis with success and was also associated with reduced episodes of infection with CoNS ((Robertson et al., 2020). More work can help develop strategies to identify reservoirs and transmission routes of NRCS-A and to eradicate carriage from neonates to prevent infection which can have serious consequences.

## Data accessibility

Sequence data for all isolates have been uploaded to the SRA under Project number PRJNA751027

## Ethics

The collection of weekly neonatal skin swabs in 2017-18 received prior approval as a surveillance study by the Research and Development Manager of the Norfolk and Norwich University Hospitals NHS Foundation Trust and did not require formal research ethics review. In Germany, the study was reviewed and approved as a surveillance study by the University of Lubeck Hospital ethics committee (reference AZ 15-034, amendment 01/2018). Clinical isolates were provided under NHS Research Ethics Committee approval to the Norwich Biorepository which banks blood, solid tissue and bacterial isolates from the NNUH and research institutes on the Norwich Research Park, including the University of East Anglia, and makes these available to the research community.

## Author Contributions

Dheeraj Sethi, Heather Felgate, Paul Clarke and Mark Webber were involved in study conception and design. Collection and processing of the Staphylococci library was carried out by Dheeraj Sethi. Kirstin Faust, Cemsid Kiy, Christoph Härtel and Jan Rupp were involved in the isolate collection from the German NICU. Heather Felgate, Dheeraj Sethi, Rebecca Clifford and Rachel Dean were involved with sequencing strains. Heather Felgate carried out phenotyping, data analysis and genotyping of the collection, as well as writing the first draft of the manuscript. Catherine Tremlett provided isolates from LOS and provided input on the manuscript. Heather Felgate, Christoph Härtel, Cemsid Ki, John Wain, Gemma Langridge, Paul Clarke, Andrew Page, Mark A Webber were all involved in the production of the manuscript and scientific input.

## Acknowledgements

We would like to acknowledge the nurses within NICUs in Lubeck and Norwich who helped with isolation of S. *capitis* from neonates and the microbiology lab staff at the NNUH and Lubeck hospital who also helped process and save isolates.

## Supplementary material

**Supplementary Data 1** List of reference genomes of *S. capitis* strains used.

**Supplementary Data 2** Meta data for *Staphylococcus capitis* isolates. MICs OCT (octenidine range 1 – 32 μg/mL), CHX (chlorhexidine range 4 - 64 μg/mL), VAN (vancomycin range 0.25 - 4 μg/mL), GEN (gentamicin range 0.016- 16 μg/mL), PEN (benzylpenicillin range 0.016 - 128 μg/mL), CEF (cefotaxime range 0.125 - 2 μg/mL), CIP (ciprofloxacin range 0.064 - 2μg/mL), DAP (daptomycin range 0.25 - 2 μg/mL), FUS (fusidic acid range 0.064 −2 μg/mL). X indicates MIC data was not obtained.

**Supplementary Data 3** ABRICATE summary in Binary format, where 1 = ≥ 90% gene identity

**Supplementary Data 4** Snippy-core SNP distance matrix on all *S. capitis* collection using NRCS-A CR05 as the reference genome.

**Supplementary Data 5** Snippy-core SNP distance matrix on NRCS-A isolates using CR05 as the reference genome.

**Supplementary Data 6** Scoary report on NRCS-A traits compared to proto NRCS-A, Country (UK), cefoxitin ≥ 4 μg/mL (CEF4), chlorhexidine ≥ 16 μg/mL (CHX 16), ciprofloxacin ≥ 2 μg/mL (CIP2), fusidic acid ≥ 2 μg/mL (FUS2), gentamicin ≥ 8 μg/mL

